# CD4 T-cell transcriptome expression reversal of the lncRNA-mRNA co-expression network in elite controller *vs.* normal-process HIV patients

**DOI:** 10.1101/606731

**Authors:** Chaoyu Chen, Xiangyun Lu, Nanping Wu

## Abstract

Elite controller refers to a patient with human imunodeficienvcy virus infection with an undetected viral load without anti-viral treatment. Studies on gene expression and regulation in these individuals are limited but significant. We enrolled 196 patients and collected CD4 T-cell samples from two elite controllers, two normal-process infected patients, and two healthy controls to perform second-generation transcriptome sequencing. Using the Cuffdiff model, we identified differentially expressed mRNAs and long non-coding RNAs with corrected P value < 0.05, and constructed a protein-protein interaction network as well a long non-coding RNA-mRNA co-expression network based on the Pearson correlation coefficient. Interestingly, some interactions within the networks were identified as associated with viral infections and immune responses. This was the first study to examine gene transcription in elite controllers and to study their functional relationships. Our results provide a reference for subsequent functional verification at the molecular or cellular level.

**Author Summary:** Some individuals can spontaneously inhibit HIV replication after infection with HIV, and thus lack any symptoms. Studies on these patients, termed elite controllers (ECs) will help researchers and clinicians to understand the interrelationship between HIV and the host. In the present study, we focused on the interactions and functional relationships between significantly differentially expressed long non-coding RNAs (lncRNAs) and mRNAs in ECs *vs*. normal-process patients (NPs). RNA-sequencing was performed for six representative samples of CD4 T-cells. Using the Pearson correlation test, an lncRNA-mRNA co-expression network was constructed. Several new regulatory relationships between transcripts were revealed that might be closely related to the ability of ECs to maintain a low viral load for long periods without anti-viral treatment. For example, lncRNA *C3orf35* was upregulated in ECs *vs*. NPs and was positively related to downregulation of *GNG2* mRNA (encoding G protein subunit gamma 2), which functions in chemokine signaling pathways and HIV-1 infection. Overall, we identified certain interesting genetic interactions that will provide information about the mechanism of host suppression of viral replication.

## Introduction

Human imunodeficienvcy virus (HIV) has been studied for more than 30 years, and research has found that HIV invades host CD4+ cells, integrates its own DNA into the host genome, and establishes a reservoir in the early stage, followed by massive replication, which destroys the normal immune system function of the host, and triggers various concurrent symptoms [1–2]. Ultimately, HIV infection can cause the host to die. Although anti retroviral therapy (ART) treatment can partially reconstitute the immune function of the host, it cannot eradicate the latent HIV pool in the host, which always maintains a low level of virus replication [3]. However, among HIV-infected individuals, a small number of patients are found to be inherently resistant to viral replication, and spontaneously induce suppression of the latent pools without any anti-viral therapy, resulting in an undetectable viral load in plasma for a long period [4–5]. These patients, termed elite controllers (ECs), have attracted significant research interest. Studying the characteristics of the autoimmune factors of ECs is anticipated to identify important factors that control virus replication. Such valuable information could lead to novel methods to treat and alleviate HIV infection. Long-noncoding RNAs (lncRNAs) are transcribed RNA molecules greater than 200 nt in length that regulate gene expression by diverse, but as yet not completely understood, mechanisms [6]. Although the function of most lncRNAs is unknown, several have been shown to regulate gene expression at multiple levels from DNA to phenotype [7]. Through a variety of means, including *cis* (near the site of lncRNA production) or *trans* (co-expressed with their target gene) mechanisms, lncRNAs play a vital role in many biological processes [8]. Studies on lncRNAs have become a hotspot in current non-coding RNA research.

Recent studies focused on the role of lncRNAs in HIV pathogenesis, especially the relationship between lncRNA regulation of gene expression and viral infection, replication, and latency [9–12]. A few lncRNAs have been characterized and proven to be closely associated HIV, for example, the knockdown of host lncRNA *NEAT1* enhanced virus production by increasing nucleus-to-cytoplasm export of Rev-dependent instability element (INS)-containing HIV-1 mRNAs [13]. The knockdown of lncRNA *NRON* enhanced HIV-1 replication through increased activity of nuclear factor of activated T-cells (NFAT) and the viral long terminal repeat (LTR) [14]. Similarily, lncRNA *MALAT1* releases epigenetic silencing of HIV-1 replication by displacing the polycomb repressive complex 2 from binding to the LTR promoter [15]. Furthermore, an HIV-encoded antisense lncRNA, *ASP-L*, was proven to promote latency HIV [16–17]. Although, the mechanism of lncRNA function is sometimes elusory and unpredicted, many current studies, which are limited to the screening and functional prediction stage based on chip or sequencing results, also provide us with valuable information and a basis for mechanistic research.

In the present study, we conducted a transcriptomics investigation for HIV elite controllers, to identify differences in the transcriptional expression profiles between elite controllers and normal-process HIV patients as compared with healthy controls. The obtained lncRNA-mRNA co-expression network revealed the possible role of the functional relationships between lncRNAs and mRNAs in the ability of elite controllers to inhibit viral replication.

## Results

### Subjects

A total of 196 individuals infected with HIV were enrolled in our cohort, including patients under treatment (179, 91.3%), untreated normal-process patients (15,7.7%) and elite controllers (2, 1.0%). In addition, we recruited two healthy individuals from the out-patient department of the hospital. Most patients were infected with the virus through heterosexual contact, followed by intravenous infection, and homosexual transmission. Based on the basic principle of intra-group identity, we determined three men and three women who were assigned to each of the three study groups as study subjects. Their ages ranged from 39 to 54 and were free of other diseases, such as tuberculosis, diabetes, or hepatitis. Specific details of the clinical characteristics of the participants are shown in S1 Table. Descriptive statistics are reported as counts (percentage) for dichotomous and categorical variables, and the median (the 25th percentile and 75th percentile) for continuous variables. The experimental design and analysis process of this research are shown in a flowchart (Fig 1).

**Fig. 1.**
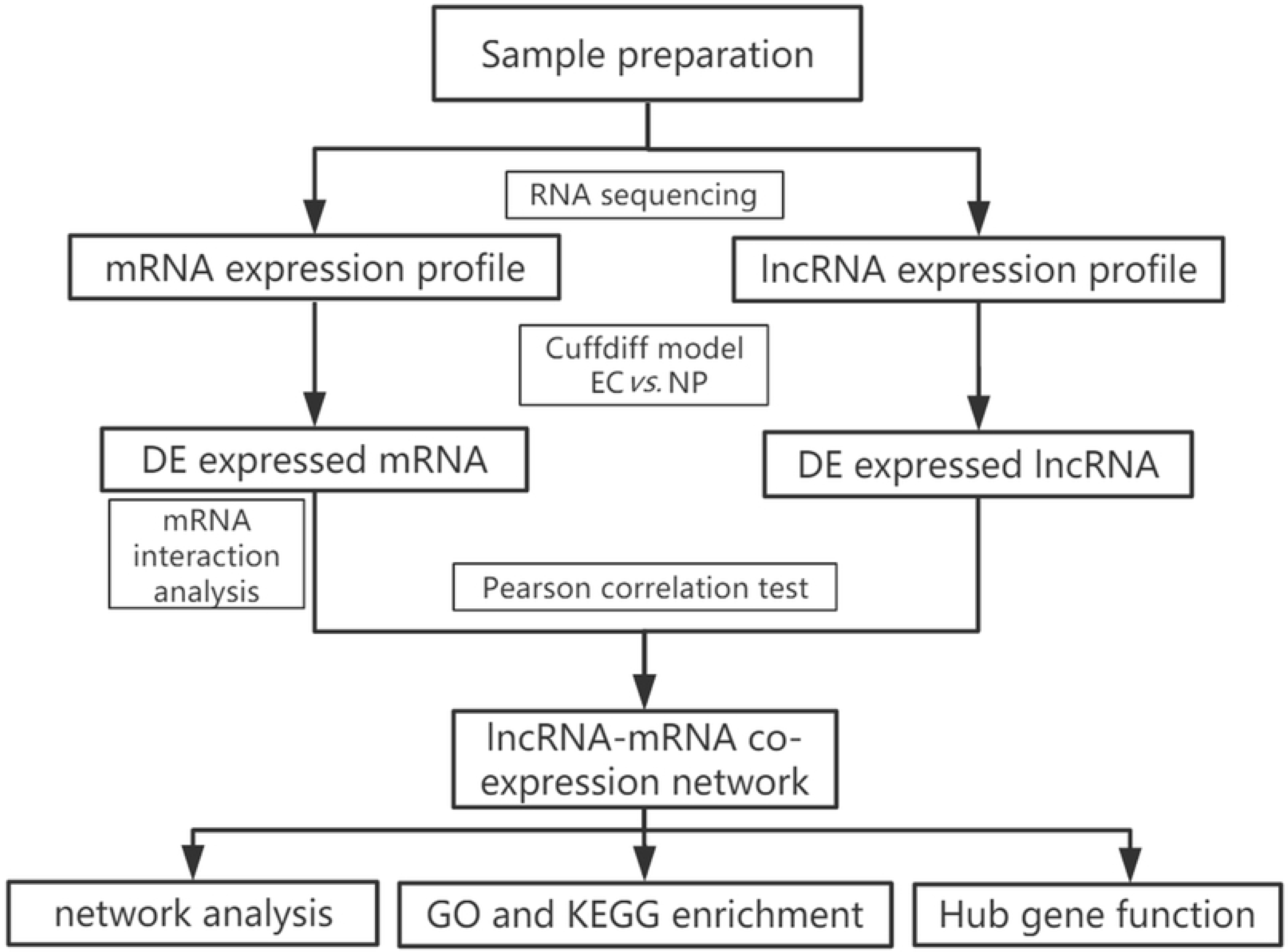
Flowchart of the analysis procedure. lncRNA, long non-coding RNA; EC, elite controller; NP, normal-process infected patient; DE, differentially expressed; GO, gene ontology; KEGG, Kyoto Encyclopedia of Genes and Genomes.

### Identification of differentially expressed (DE) transcripts between health controls (HCs) and normal-process infected patients (NPs)

First, we scanned the expression profiles of all transcripts in CD4 T-cells from two HCs, two ECs and two NPs. In all, more than 82000 mRNA transcripts and more than 20000 corresponding genes were identified. Likewise nearly 15000 lncRNAs and 1000 corresponding gene IDs were identified. Regular grouping between HCs and normal-process patients was first considered. Altoghter, 3602 mRNA transcripts and 383 lncRNA transcripts were differently expressed according to the threshold of fold change > 2 and p value < 0.05, among which 2313 mRNAs and 207 lncRNAs were upregulated and 1289 mRNAs and 176 lncRNAs were downregulated. We similarly compared gene expression differences between HCs and ECs though more attention was paid to ECs *vs.* NPs. The comparation with HCs helped to provide a basic reference background. The lists of fragments per kilobase of transcript per million mapped reads (FPKM) values and regulatory situation of all DE mRNAs and lncRNAs in HCs *vs.* NPs and HCs *vs.* ECs are shown in S2 Table and S3 Table respectively.

### Identification of DE mRNAs between ECs and NPs

Among all transcripts, 2936 transcripts were differentially expressed according to the threshold of fold change > 2 and p value < 0.05. There were 1327 mRNAs that were upregulated and 1609 that downregulated in ECs *vs*. NPs. Gene ontology (GO) and Kyoto Encyclopedia of Genes and Genomes (KEGG) pathways analysis was carried out to annotate the function of these DE mRNAs. Certain terms and pathways that might be associated with infection and immunity were significantly enriched (S1 Fig). These terms and pathways included negative regulation of interferon-beta production (GO:0032688), cellular component disassembly involved in execution phase of apoptosis (GO:0006921), regulation of defense response to virus by virus (GO:0050690), negative regulation of type I interferon production (GO:0032480), RIG-I-like receptor signaling pathway (hsa04622), and antigen processing and presentation (hsa04612).

### Identification of DE lncRNAs between ECs and NPs

As same, among all transcripts, 3543 official gene symbols that matched in the NONCODE database had corresponding lncRNA transcripts. Three main categories of lncRNAs accounted for the majority, including antisense lncRNAs (2537/14853, 17.1%), intronic lncRNAs (8152/14853, 54.9%), and long intergenic noncoding RNAs (lincRNAs; 3344/14853, 22.5%). There were 151 upregulated lncRNAs and 160 downregulated lncRNAs in ECs *vs*. NPs according to the threshold of fold change > 2 and p < 0.05. In both the mRNA and lcnRNA clusters (Fig 2), the samples showed marked intra-group correlations and inter-group differences, which further confirmed the specificity of the samples. In terms of protein-coding mRNAs encoding proteins with specific biological functions, the HC and EC groups showed closer clustering and were relatively distant from the NP group. The list of fragments per kilobase of transcript per million mapped reads (FPKM) values and regulatory situation of all DE mRNAs and lncRNAs are shown in S4 Table.

**Fig. 2.**
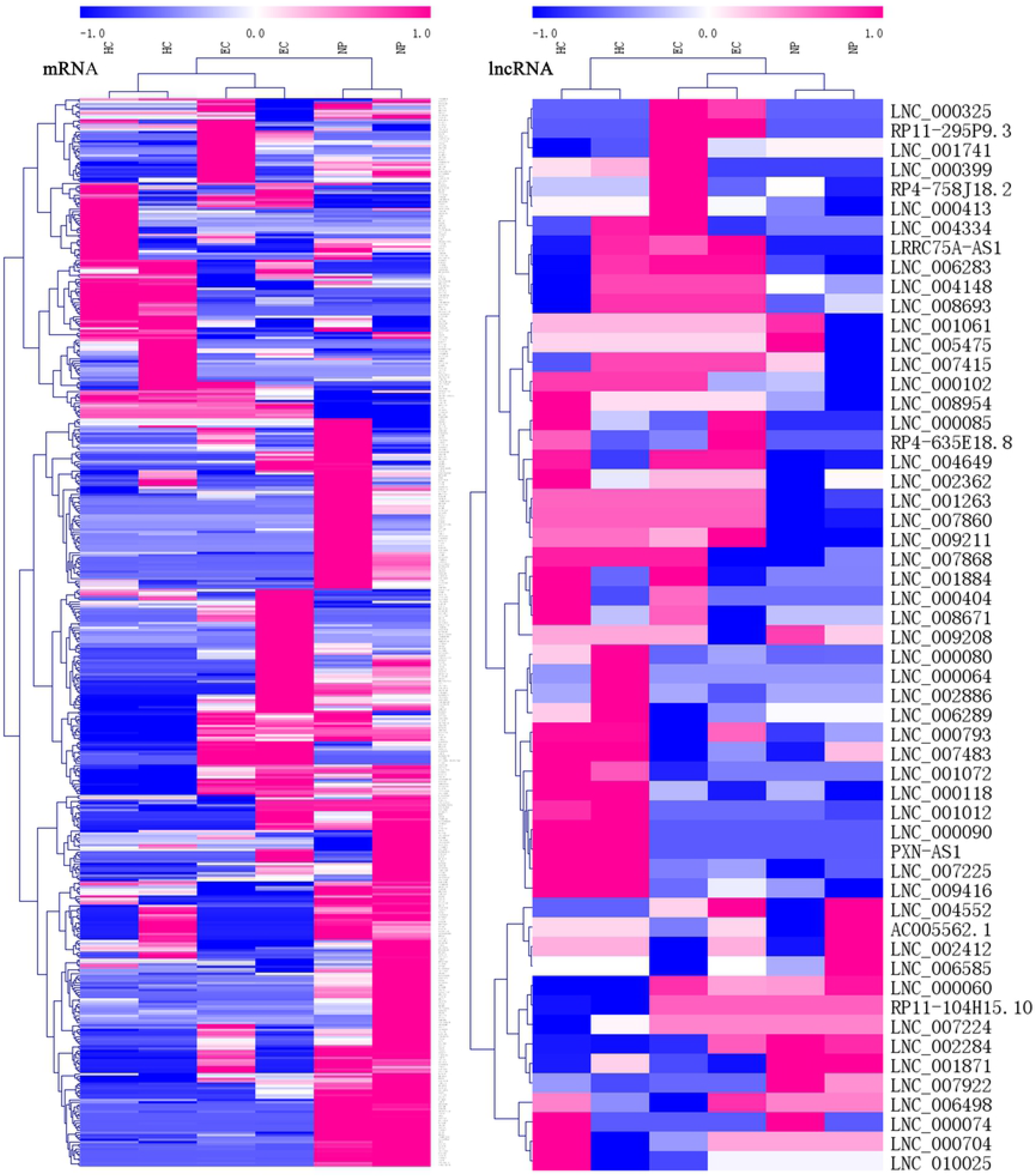
Heatmaps of differentially expressed (DE) mRNAs and long non-coding RNAs (lncRNAs). Hierarchical clustering analysis of sequencing detected mRNAs and lncRNAs with their expression abundance. Patient groups are on the horizontal axis, and mRNA and lncRNA genes are grouped along the vertical axis. Pinkish-purple bars indicate increased abundance of the corresponding genes, and blue indicates decreased abundance. White bars indicate that the corresponding genes were not detected.

### PPI network of DE mRNAs

Fig 3 shows the top 50 hub genes playing an important role in interaction of DE mRNAs, in which many genes functioned centrally and could be frequently regulated by other transcripts. Considering the differences in the outputs of different algorithms, we identified those genes that were commonly detected by the algorithms and focused on those genes that were regulated by lncRNAs and their functional enrichment analysis. S5 Table shows the intersection list of the top 50 hub-mRNAs calculated using 12 algorithms.

**Fig. 3.**
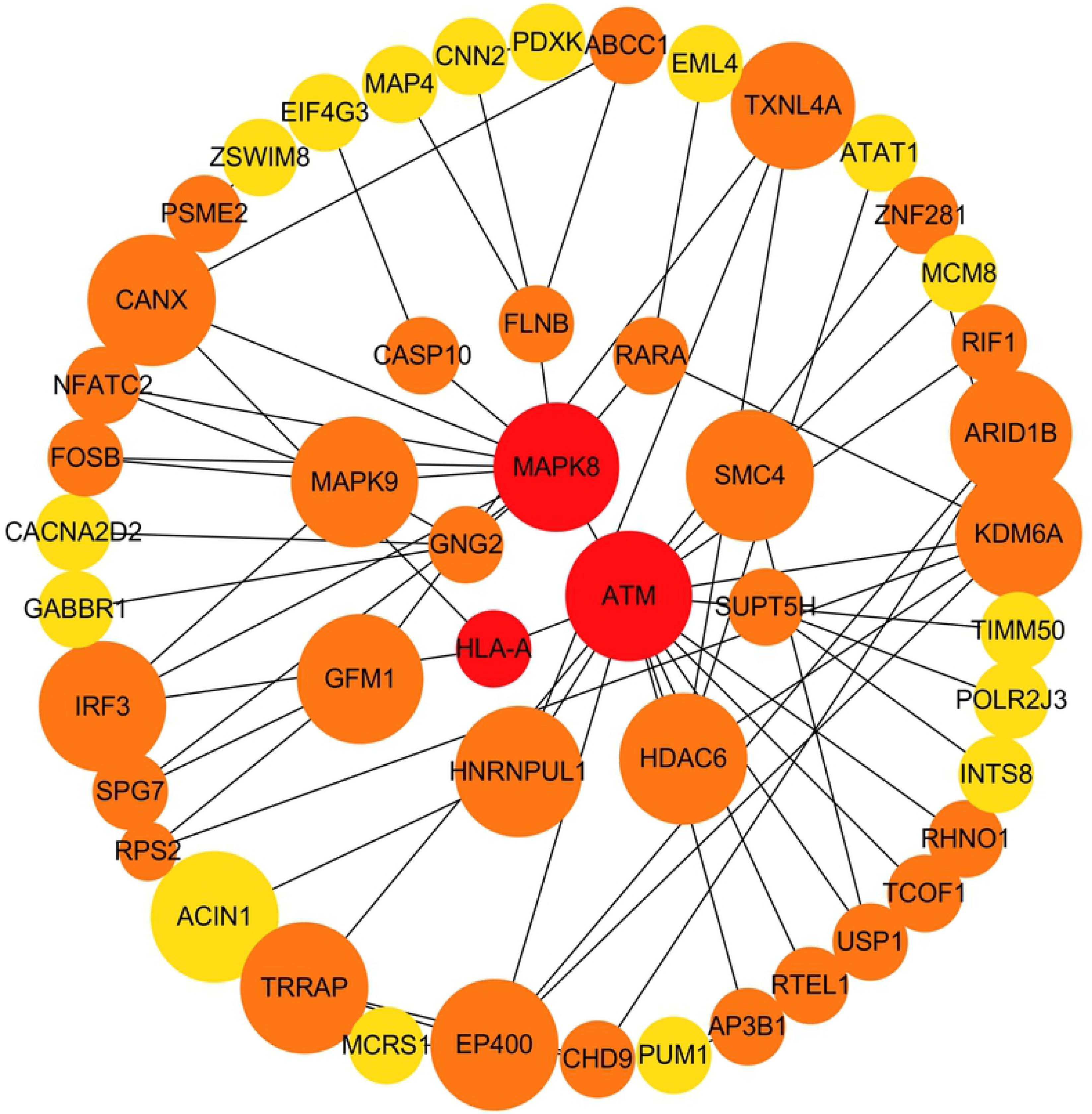
Protein-protein interaction (PPI) network of differentially expressed (DE) mRNAs. Top 50 hub mRNAs were included in using EcCentricity algorithm. The intensity of the color indicates the density of the correlation. The closer to red, the more central the gene is. The size of the circle represents the significance of a gene.

### LncRNA-mRNA co-expression network analysis

Almost 0.9 million co-expression relationships were discovered, and the exact form was expressed as a linear orientation whereby one lncRNA targets one mRNA. In all orientations, 1109 DE lncRNAs targeting DE mRNAs were identified. Fig 4a shows the network constructed using these relationships, in which 142 DE lncRNAs were co-expressed with 127 DE mRNAs. Moreover, we found 16 of 24 intersecting hub mRNAs in the diagram, which helped us to quickly locate the target protein coding genes and easily access the enriched functions of these genes. Next, we selected 15 annotated lncRNAs and their associated mRNAs from Fig 4a and constructed Fig 4b. We then identified the top 10 GO terms and pathways most closely related to infection and immunity by combining the function of the hub genes. Many significant functions are co-enriched, such as immune response-regulating signaling pathway (GO:0002764) and apoptosis (hsa04215) (Fig 5).

**Fig. 4.**
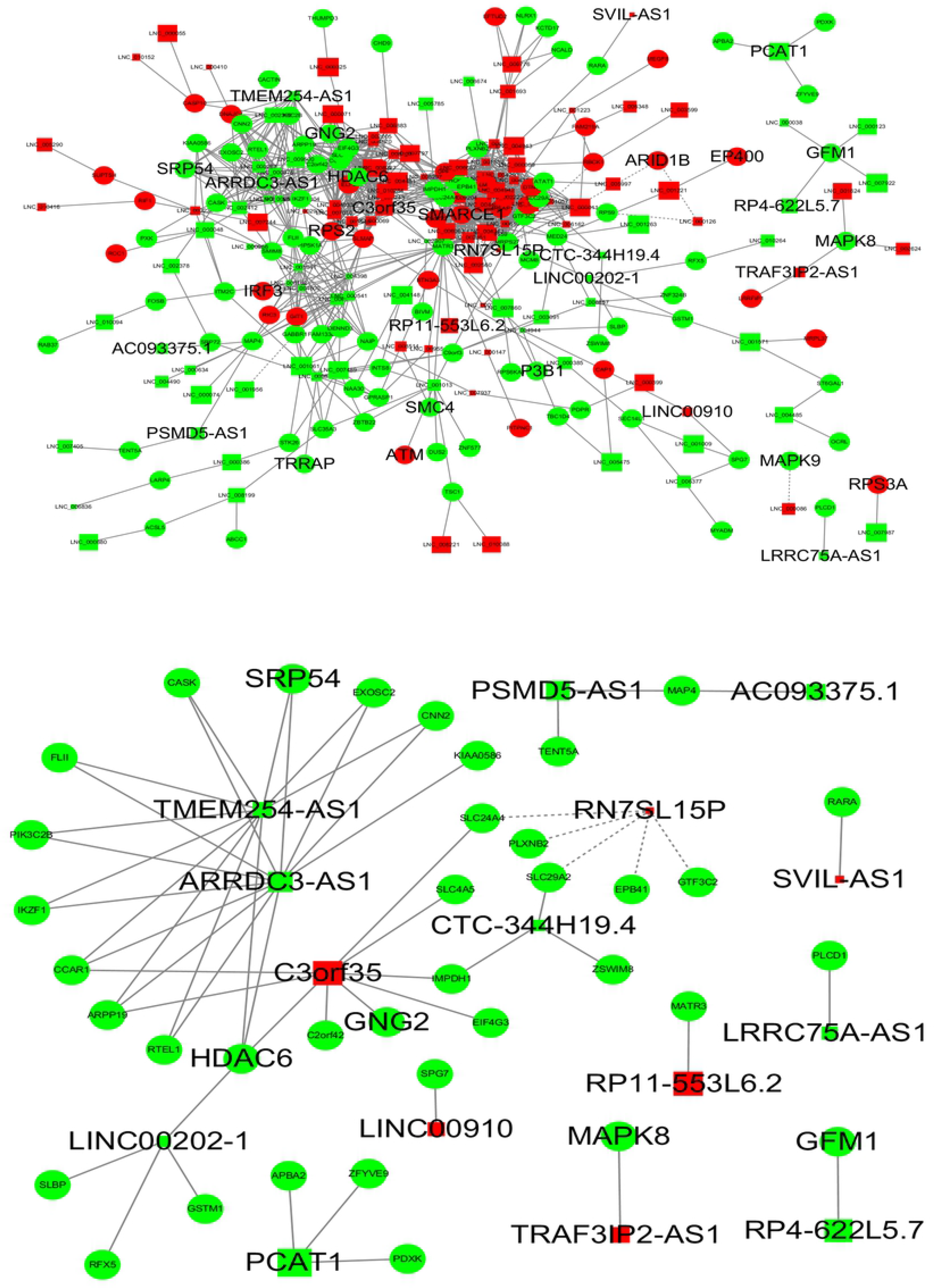
Long non-coding RNA (lncRNA)-mRNA co-expression network in elite controllers (ECs) and normal-process infected patients (NPs). The network was based on the Pearson relation coefficient of an lncRNA targeting an mRNA. Round shapes correspond to mRNAs, and squares to lncRNAs. The size of the shape is positively related to the P value. Red means upregulation and green means downregulation. A solid line indicates a positive correlation and a dotted line indicates a negative correlation. All hub-mRNAs and annotated lncRNAs are tagged with large labels. (a) All lncRNAs and co-expressed mRNAs. (b) Annotated lncRNAs and co-expressed mRNAs.

**Fig. 5.**
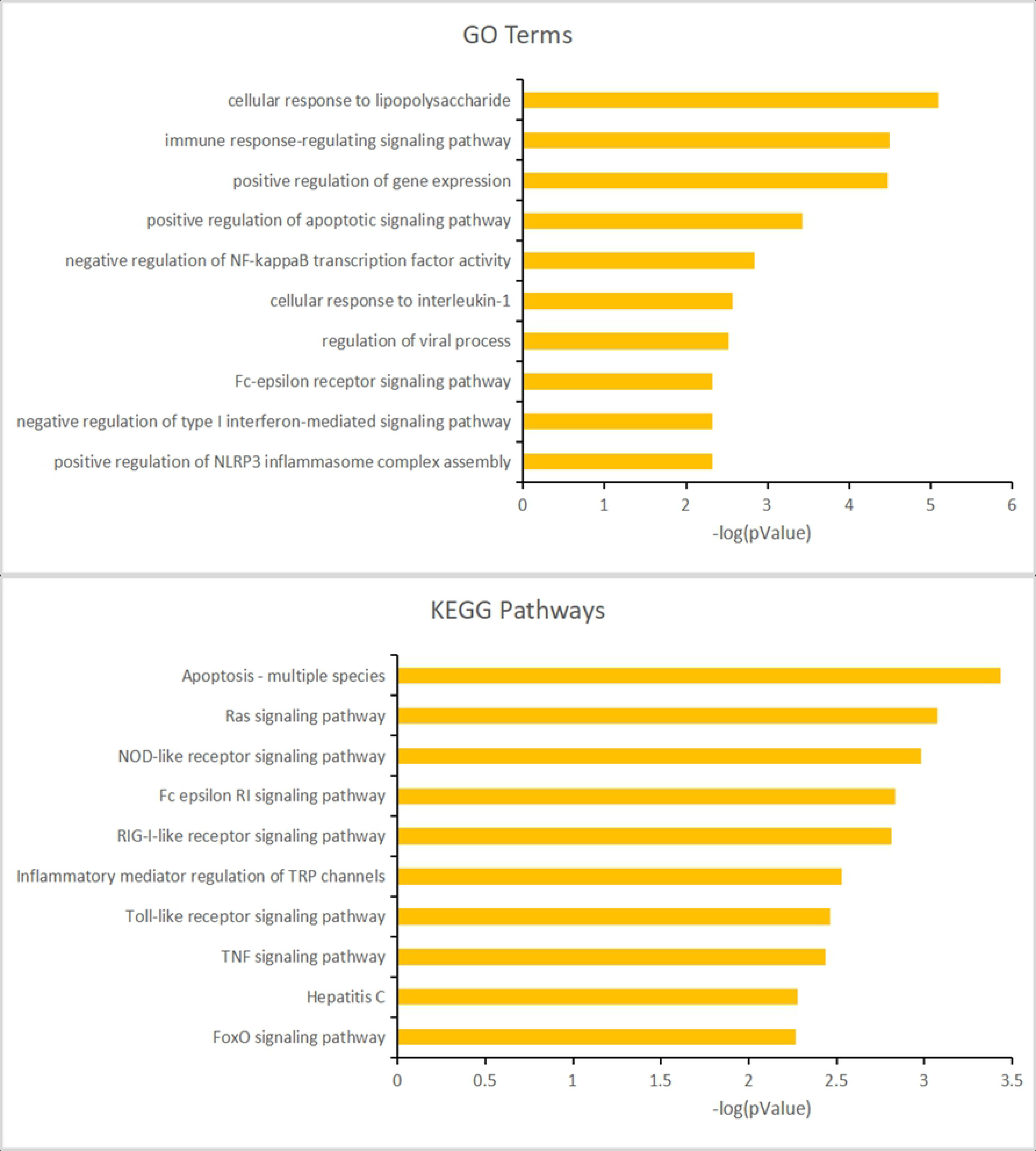
Gene ontology (GO) terms and Kyoto Encyclopedia of Genes and Genomes (KEGG) enrichment classification of the predicted targeting mRNAs of the long non-coding RNA(lncRNA)-mRNA network. The 10 most significantly enriched KEGG pathways and the 10 most significantly enriched GO terms in the biological process, cellular components, and molecular function categories.

### Immune regulation and infection associated lncRNA-mRNA networks

We further investigated the specific functions of the co-expressed mRNAs shown in Fig 4b and found several interesting regulation relationships. Certain genes were enriched to GO terms and pathways that are closely related to immune regulation and infection. These genes and corresponding pathways included *GNG2* (Human cytomegalovirus infection, hsa05163; Chemokine signaling pathway, hsa04062; Kaposi sarcoma-associated herpesvirus infection, hsa05167; Human immunodeficiency virus 1 infection, hsa05170), *RFX5* (Primary immunodeficiency, hsa05340; Tuberculosis, hsa05152; Antigen processing and presentation, hsa04612), *HDAC6* (Viral carcinogenesis, hsa05203), *MAPK8* (Apoptosis - multiple species, hsa04215; NOD-like receptor signaling pathway, hsa04621; Fc epsilon RI signaling pathway, hsa04664; RIG-I-like receptor signaling pathway, hsa04622; Inflammatory mediator regulation of TRP channels, hsa04750; Toll-like receptor signaling pathway, hsa04620), *MAPK4* (IL-17 signaling pathway, hsa04657), and *RARA* (Th17 cell differentiation, hsa04659). Corresponding lncRNAs including *C3orf35*, *TMEM254-AS1*, *ARRDC3-AS1*, *LINC00202-1*, *TRAF3IP2-AS1*, *PSMD5-AS1*, *AC093375.1*, and *SVIL-AS1* were co-expressed with these mRNAs, which suggested several new regulatory relationships that might play an important role in controlling virus replication.

## Dicussion

In the present study, we systematically analyzed the expression profiles of lncRNAs and mRNAs from CD4+ T cells from ECs, NPs, and HCs, and constructed a co-expression network based on the relationships among DE genes and database annotations. The functions of most lncRNAs are not well characterized; therefore, the gene functions in this research mainly referred to those of mRNAs and their expression products. The potential functions of candidate lncRNAs and the lncRNAs-associated biological processes in ECs were predicted using the lncRNA-mRNA network and functional enrichment analysis. We identified certain genes that play a significant role in the ability of ECs to inhibit the replication of latent virus. Most of these genes were downregulated in the HCs and ECs and upregulated in the NPs, which was consistent with their clinical characteristics, in that ECs maintain a similar gene expression pattern to the HCs, which distinguishes them from the NPs. Among these rigorously selected genes, some hub genes were closely related to microbial infections and host immune processes, which are reported to greatly affect the progress of HIV infection [18–19].

To the best of our knowledge, the present study was the first to predict lncRNA-mRNA interactions for HIV ECs, in which many of the reported genes had not been previously reported to function in HIV-associated biological process. The *MAPK* family has been associated with the reactivation and replication of the virus in many studies [20–21], and the HIV-1 virus inhibition activity of MAPK pathway inhibitors was the result of the negative regulation of HIV-1 LTR promoter activity [22]. In addition, HDAC6 gene expression product was reported to mediate HIV-1 tat-induced proinflammatory responses by regulating the MAPK-NF-kappaB/AP-1 pathways in astrocytes [23]. In the present study, we also found that HDAC6 interacts with MAPK8, MAPK4, and MAPK9 (Fig 3). This suggested their related lncRNAs including *C3orf35*, *TMEM254-AS1*, *ARRDC3-AS1*, *LINC00202-1*, *TRAF3IP2-AS1*, *PSMD5-AS1*, and *AC093375.1* may control HIV replication by regulating the expression of these genes. In addition, we identified genes that had not been studied before, whose functions were enriched into the following processes: Virus infection, immunodeficiency, apoptosis, and cytokines. Previous studies had shown that these functions are closely related to viral replication [24–25]. For instance, *GNG2* (encoding G protein subunit gamma 2) showed a strong correlation with HIV in the KEGG pathway analysis. In Kaposi sarcoma-associated herpesvirus infection, *GNG2* activates Ras pathways and PI3K-Akt pathways to affect the production of cellular cytokines such as IL-6, IL-8, VEGF, and COX2, and other inflammatory and cellular life cycle-related biological processes. HIV infection and its concurrent Kaposi sarcoma are also closely related to the occurrence of other cancers [26–27].

ART treatment cannot eradicate the latent virus in the host; therefore, the genes identified in the present study may play a non-negligible regulatory role in the relationship between the latent virus and the host. Latent virus is the biggest obstacle to the elimination of the disease; thus, understanding the processes that contribute to its persistence, such as inflammation and immune activation, are crucial for the remission and cure of HIV [28–29].

There are also some limitations to this study. First, because of the limited amount of RNA in the samples, the sample size of this study was relatively small. Although we selected eight genes for verification using qRT-PCR in 12 samples and found that the expression trends of four of them were consistent with the RNA sequencing results, these results are not statistically significant because of the small sample size. Second, we only assessed gene expression of CD4+ T cells, which, although they are the main targets of HIV, are only one of the many cell types that are infected by HIV. Further research should be performed to validate the function and mechanism of the genes in a larger sample and in more cell types.

In summary, this study identified several important differentially expressed genes associated with the EC phenomenon, which could form the basis for subsequent cellular and molecular studies, and could provide new targets for gene-targeted therapy in the future.

## Methods

### Ethics statement

All participating medical institutions provided local institutional review board approval, and the participants provided informed consent for this study.

### Cohort and patients

In total, 196 patients were recruited into our cohort, from different provinces of China, who were diagnosed in Qingchun Hospital of Zhejiang Province from January 2000 to January 2018, and received or voluntarily rejected highly active antiretroviral therapy (HAART) treatment. The viral loads of the patients were detected using the Cobas Taqman system (Roche). The HCs were randomly selected from hospital clinics and were not suffering from any diseases. We chose two ECs, two NPs, and two HCs of similar ages, gender, and traditional risk factors to perform second generation transcriptome sequencing. ECs were defined as HIV-infected patients with and undetectable viral load without any anti-viral treatment for nearly 10 years.

### Sample collection and preparation

Peripheral blood mononuclear cells (PBMCs) were isolated from 3–5 mL EDTA-K2+ anticoagulant venous blood using density gradient centrifugation. Primary CD4+ T cells were purified through negative selection using a CD4+ T cell isolation Kit (Miltenyi). According to flow cytometry analysis, the purity of the separated CD4+ T cells was more than 90%. Total RNA was extracted using the TRIzol reagent (Life Technologies). RNA purity was checked using a NanoPhotometer^®^ spectrophotometer (IMPLEN). The RNA concentration was measured using a Qubit^®^ RNA Assay Kit in Qubit^®^ 2.0 Fluorometer (Life Technologies) and its integrity was assessed using an RNA Nano 6000 Assay Kit for the Bioanalyzer 2100 system (Agilent Technologies).

### RNA library construction and RNA sequencing

Ribosomal RNA was removed using a Epicentre Ribo-zero™ rRNA Removal Kit (Epicentre), and the rRNA free residue was cleaned up using ethanol precipitation. Subsequently, sequencing libraries were generated using the rRNA depleted RNA by NEBNext^®^ Ultra™ Directional RNA Library Prep Kit for Illumina^®^ (NEB) following manufacturer’s recommendations. After cDNA synthesis and purification, clustering of the index-coded samples was performed on a cBot Cluster Generation System using a TruSeq PE Cluster Kit v3-cBot-HS (Illumina) according to the manufacturer’s instructions. The libraries were then sequenced on an Illumina Hiseq 4000 platform and raw data were generated.

### RNA sequencing analysis

Clean data were obtained by removing reads containing adapters, reads only containing poly-N, and low quality reads from the raw data. After quality control, paired-end clean reads were aligned to the reference genome downloaded from website using HISAT2 (v2.0.4). Cuffdiff (v2.1.1) was used to calculate the FPKM values of both lncRNAs and mRNAs in each sample [30]. Gene FPKM values were computed by summing the FPKM values of transcripts in each gene group. The Cuffdiff model provides statistical routines to determine differential expression in the digital transcript or gene expression data using a model based on the negative binomial distribution. Transcripts with a corrected P value < 0.05 were assigned as differentially expressed. We plotted a heatmap to observe the clustering between the samples and the genes using Multi Experiment Viewer (v4.9.0).

### LncRNA-mRNA co-expression network analysis

*Trans* role is lncRNA binding to target DNA as a RNA:DNA heteroduplex, as RNA:DNA:DNA triplex, or RNA recognition of specific chromatin-like complex surfaces[31]. We calculated the expression correlation between lncRNAs and mRNAs using the Pearson correlation test for target gene prediction and the results were expressed as Pearson correlation coefficients. To narrow the scope of investigation and the number of genes, we first assessed the interaction among all DE mRNAs using the STRING database (v11.0) and constructed a PPI network using the Hubba package in Cytoscape (v3.7.1). After considering 12 synthetic algorithms, the intersection of the top 50 genes yielded 24 hub genes, which occupy a central position in the network of all DE mRNAs. After identifying and retaining the DE mRNAs and lncRNAs in all lncRNA-mRNA relations, we screened 1109 pairs of exact relationships between the two groups and constructed a co-expression network based on the Pearson relation coefficient (all > 0.95). Then, we identified the annotated lncRNAs and associated mRNAs for further functional enrichment analysis. PPI and lncRNA-mRNA network diagrams were also drawn using the Cytoscape software.

### Gene ontology and KEGG pathways analysis

By jointly using the David database (v6.8) and KOBAS database (v3.0), gene GO enrichment and KEGG pathways analysis of all DE mRNAs and DE lncRNA co-expressed DE mRNAs were implemented to interpret the biological meaning of the transcripts. GO terms and KEGG pathways with corrected P values less than 0.05 were considered significantly enriched by the differentially expressed RNAs.

## Acknowledgements

We would like to thank all of the participants in the cohort. In addition, we would like to thank the native English speaking scientists of Elixigen Company (Huntington Beach, California) for editing our manuscript.

## Author Contributions

Chaoyu Chen and Nanping Wu were involved in the conception and design of the study. Chaoyu Chen and Xiangyun Lu were involved in sample preparation, data acquisition, and organization. Chaoyu Chen conducted the data analysis, and all authors were responsible for data interpretation. Chaoyu Chen was the major contributor to the writing of the manuscript. All authors read and approved the final manuscript.

## Supporting information

**S1 Figure. Gene ontology (GO) terms and Kyoto Encyclopedia of Genes and Genomes (KEGG) pathways enrichment of all differentially expressed (DE) mRNAs.**

**S1 Table. Clinical characteristics of the participants.**

**S2 Table. Lists of differentially expressed (DE) mRNAs and long non-coding RNAs (lncRNAs) in health controls (HCs) *vs*. normal-process infected patients (NPs).**

**S3 Table. Lists of DE mRNAs and lncRNAs in HCs *vs*. elite controllers (ECs).**

**S4 Table. Lists of DE mRNAs and lncRNAs in ECs *vs*. NPs.**

**S5 Table. The intersection list of the top 50 hub-mRNAs calculated using 12 algorithms.**

